# Mapping adaptive immune responses toward fungal antigens in inflammatory bowel disease using T cell repertoire sequencing and phage-immunoprecipitation sequencing

**DOI:** 10.1101/2025.05.13.653535

**Authors:** Aya K.H. Mahdy, Hesham ElAbd, Melanie Prinzensteiner, Hannah Jebens, Kostas Sivickis, Petra Bacher, Thomas Vogl, Mathilde Poyet, Andre Franke

## Abstract

Inflammatory bowel disease (IBD) is an idiopathic, immune-mediated chronic inflammatory disease of the gut with two primary clinical forms, Crohn’s disease (CD) and ulcerative colitis (UC). Several genetic susceptibility variants have been associated with IBD, such as *ATG16L, NOD2,* and several human leukocyte antigen (HLA) alleles, nonetheless, the actual disease causes remain unknown. Whereas previous findings have shown elevated responses toward fungal antigens in individuals with IBD, *e.g.,* elevated anti–*Saccharomyces cerevisiae* antibody (ASCA) levels, an exhaustive mapping of immune responses toward fungal antigens remains incomplete. Thus, we analyzed the fungal mycobiome profiled using internal transcribed spacer 2 (ITS2)-sequencing simultaneously with the T cell repertoire of 637 individuals with IBD from the SPARC IBD cohort, which enabled us to identify 31 T cell clonotypes targeting several prevalent members of the gut mycobiome. Subsequently, we developed a novel phage-immunoprecipitation sequencing (PhIP-Seq) library covering 12,000 potential antigens from the proteome of *S. cerevisiae* and screened for antibody responses in 100 individuals with CD and 60 healthy controls with known ASCA status, enabling us to identify public and private antibody responses against several *S. cerevisiae* proteins. In conclusion, we corroborated previous findings showing elevated T cell responses against fungal antigens in individuals with IBD and identified multiple antigenic proteins from the proteome of *S. cerevisiae* that are targeted by the immune system of individuals with and without CD.

## Introduction

IBD is an immune-mediated inflammatory disease of the gut with multiple genetic and environmental risk factors^1–6^. While the exact etiology remains unknown, several aberrant immune responses have been identified in individuals with IBD, such as dysregulated responses toward the gut microbiota^7^ and increased T helper 1 (Th1) responses toward prevalent members of the gut mycobiome, such as *S. cerevisiae* and *C. albicans^8^*. Individuals with Crohn’s disease (CD), which is a subtype of IBD, have been shown to have a higher titer of anti-*saccharomyces cerevisiae* antibodies (ASCA)^9^ targeting mannoproteins expressed at the fungal cell wall^10^. Several studies have shown different alterations in the composition of the gut mycobiome in individuals with IBD^10–12^, nonetheless, the immunological consequences of these alterations remain poorly understood.

The last decade has witnessed the maturation of multiple high-throughput profiling assays such as T cell repertoire sequencing (TCR-Seq)^13^ and Phage-immunoprecipitation sequencing (PhIP-Seq)^14^. TCR-Seq enables the collection of T cell receptor chains present in a sample to be sequenced, *e.g.,* the α (TRA) or the β (TRB) repertoire. While TCR-Seq does not reveal the antigenic specificity of the identified TRA or TRB chains, *i.e.,* clonotypes, large-scale association studies across hundreds of individuals have identified hundreds of clonotypes that are associated with different antigenic exposures, *e.g.,* cytomegalovirus (CMV)^15^, Lyme disease^16^ and SARS- CoV-2^17^. Besides infectious diseases, large-scale TCR-Seq has been used to identify exact clonotypes involved in different immune-mediated inflammatory diseases such as IBD^18–20^ and type 1 diabetes^21^.

On the contrary, phage-immunoprecipitation sequencing (PhIP-Seq) reveals exact antigens bound by the antibodies in the sera of hundreds to thousands of individuals but without identifying the exact antibodies’ DNA sequences involved in binding these antigens. Conceptually, PhIP-Seq is composite of several steps^22^. First, designing a library of peptide antigens and *in silico* reverse translating these peptide sequences into DNA sequences that can be synthesized in a high throughput manner. Subsequently, these antigens are cloned in-frame into a suitable bacteriophage, *e.g.* the T7 bacteriophage, then, these bacteriophages are incubated with sera from the study participants. After that, bacteriophages displaying antigens recognized by the antibodies in the serum of a specific samples will be pulled down using an immunoprecipitation process that targets a specific group of antibodies, *e.g.* IgG. Hence, any unbound bacteriophage will be washed away. Lastly, the cloned DNA sequences of antibody-bound bacteriophages are amplified using PCR and sequenced to identify antigens bound the antibodies in the sera of a given sample.

PhIP-Seq has been used to identify public antigenic-exposure histories at the population level^14,23^, to map immune responses across different infections^24^, and to identify antigens involved in the pathogenesis of different diseases^25,26^. Utilizing both technologies, *i.e.* TCR-Seq and PhIP-Seq, we aim in the current study to decode the T cell responses and antigenic targets of the human mycobiome, particularly in individuals with IBD.

## Results

### Individuals with CD have elevated T cell responses against dominant gut mycobiota

Expanded anti-fungal responses in individuals with CD^8^, particularly cross-reactive clonotypes recognizing *C. albicans, C. tropicalis, and S. cerevisiae,* have been previously identified. We first aimed to validate the identified signal using a large cohort of individuals with CD and UC. Thus, we compared the expansion of fungal-specific clonotypes in the “*A Study of a Prospective Adult Research Cohort with Inflammatory Bowel Disease (SPARC IBD)*” cohort of the Crohn’s & Colitis Foundation^27^. This cohort contains 1,890 individuals with CD and 914 individuals with UC, along with detailed clinical records. The expansion of fungal-specific TRB clonotypes identified previously was higher in individuals with CD relative to UC (**Fig. 1A**). This was observed across the three fungal species previously investigated, namely, *C. albicans, C. tropicalis, and S. cerevisiae* (**Fig. 1B**). Besides disease status, we also observed a significant interaction with biological sex where anti-*S. cerevisiae* responses were significantly higher in females with CD (P=0.048; **Fig. S1A**), while anti-*C. tropicalis* responses in females with UC were significantly higher (P=0.0084; **Fig. S1B**).

**Figure 1:**
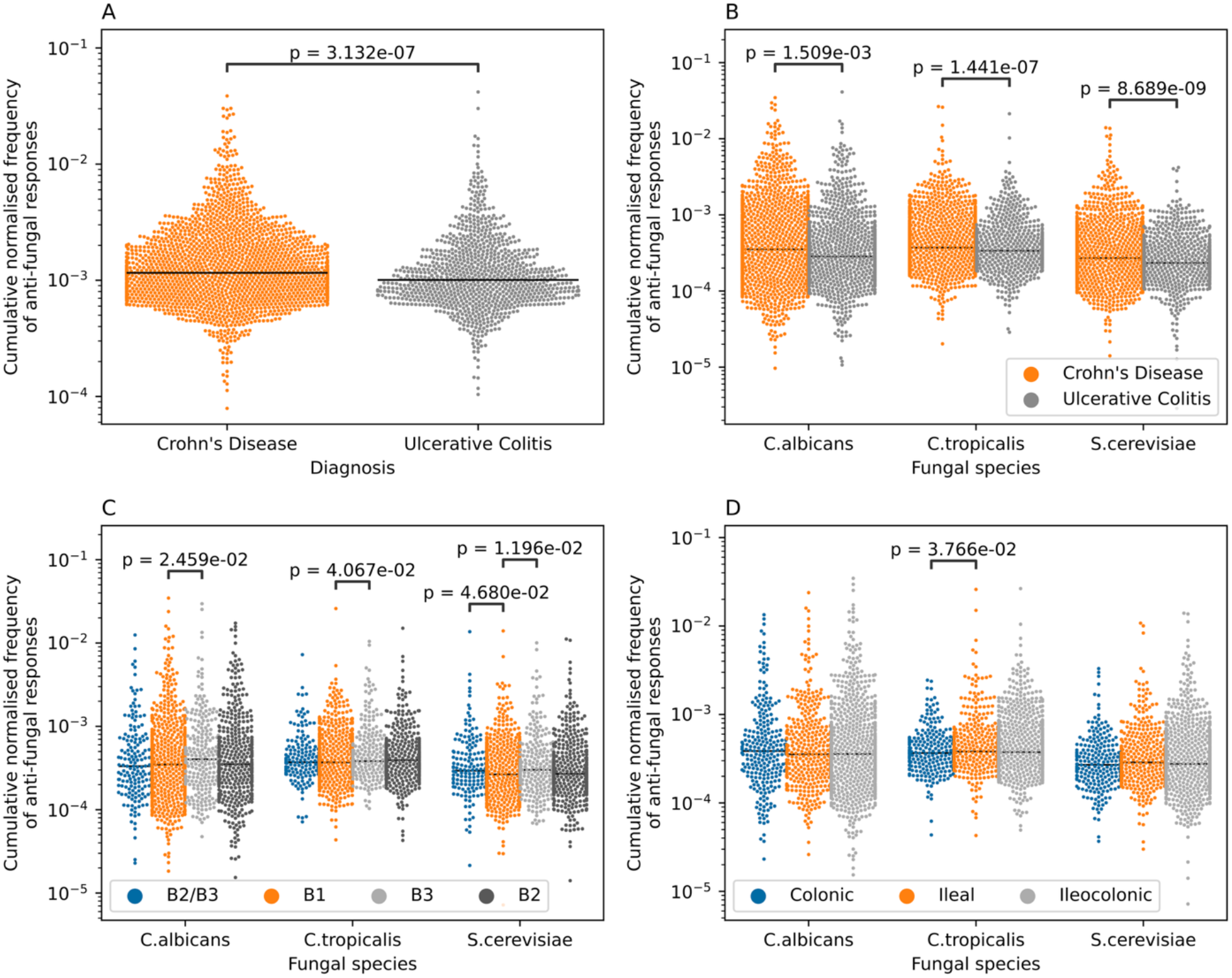
Overview of anti-fungal responses using 2,804 individuals from the SPARC IBD cohort. (**A**) shows the expansion of anti-fungal clonotypes derived from Martini *et al^8^* in 1,890 individuals with CD and 914 individuals with UC from the SPARC IBD cohort. (**B**) the expansion of anti-fungal clonotypes in individuals with CD is observed across the three fungal species, with the highest magnitude observed in responses targeting *S. cerevisiae*. (**C**) shows the expansion of diMerent anti-fungal clonotypes in individuals suMering from CD with diMerent disease behaviors (Montreal classifications), namely, B1 (non- stricturing, non-penetrating), B2 (stricturing), B3 (penetrating), and B2/B3 (stricturing and penetrating) disease. (**D**) shows the expansion of anti-fungal clonotypes in individuals with CD from the SPARC IBD cohort with diMerent disease locations, namely, ileal, colonic, and ileocolonic. Across all the panels, a two-sided Mann-Whitney Wilcoxon U test was used to compare between the diMerent groups, with only significant (P<0.05) results shown. Additionally, individuals where the expansion of these clonotypes was not detected, *i.e.,* individuals with a cumulative individual if zero, were filtered out prior to visualization and statistical comparisons.

Given the significant expansion of anti-fungal T cell responses in individuals with CD, we focused on identifying factors associated with higher expansion in these individuals. There was neither a significant association between the expansion of anti-fungal responses and age (**Fig. S2**) nor years since diagnosis (**Fig. S3**). However, the expansion was higher in individuals with a penetrating or a stricturing and penetrating disease behavior (**Fig. 1C**). Furthermore, the expansion poorly correlated with disease location, where only anti-*C. tropicalis* responses were higher in individuals with ileal CD (**Fig. 1D**). Thus, these observations corroborate previous findings^8^, showing elevated anti-fungal responses in individuals with CD.

### Differential presentation by fungal antigens in individuals with CD relative to healthy controls

To gain more insights about these clonotypes we aimed at studying their HLA restriction using our previously published dataset of HLA alleles and their restricted TRB clonotypes^28^. Out of the 18, 303 unique TRB clonotypes derived from three individuals with CD and three healthy controls, we were able to resolve the HLA restriction of 807 unique clonotypes. Out of which 584 (72.3%) were specific to one allele and 223 (27.7%) were associated to multiple alleles, mostly, residing on the same HLA haplotype due to the linkage-disequilibrium among these HLA alleles. Using clonotypes derived from individuals with CD and controls, we observed that most of these clonotypes were restricted to the HLA-DRB1*04:01 (**Fig. S4A**). By focusing on clonotypes derived from CD we observed that they were mainly restricted to the HLA-DQA1*01:01-DQB1*05:01 (**Fig. S4B**), meanwhile, the clonotypes derived from healthy controls were mainly restricted to HLA- DRB1*04:01 (**Fig. S4C**). This hints toward a differential presentation of fungal antigens in individuals with CD and healthy individuals. These findings suggest a differential presentation of fungal antigens in individuals with CD and healthy controls. Nonetheless, given the small sample size of cases and controls included in the Martini *et al.^8^* dataset (n=6 individuals) and the high HLA polymorphism, this difference could also be attributed to differences in the genetic background of the study participants. Thus, identifying fungal specific clonotypes in a larger cohort of individuals is needed to resolve and pinpoint differences in fungal antigen presentations.

### Mapping the interaction between the gut mycobiome and the TRB repertoire using statistical co-occurrence analysis

Motivated by these findings, we aimed to map the interactions between the entire gut mycobiome and the host TRB repertoire by analyzing their statistical co-occurrence patterns. Briefly, for 637 individuals from the SPARC IBD cohort^27^, the gut mycobiome was profiled using ITS2-seq in addition to their TRB repertoire (**Methods**), subsequently, we used a variant of the analytical approach developed by Emerson *et al.^15^* to identify clonotypes associated with prevalent gut mycobiota (**Methods**; **Fig. 2A**). Nonetheless, most members of the gut mycobiome were private and were not shared among different individuals with only 35 amplicon sequence variants (ASV) out of 3,036 ASV identified from the study cohort being shared among 5% or more of the study cohort (**Fig. 2B**). Using a stringent Bonferroni-based multiple testing correction threshold of P<2.8×10^−6^ only 31 TRB clonotypes were significantly associated with these 35 ASVs. By resolving the taxonomic correspondence of these ASVs we observed that the associated TRB clonotypes were targeting different members of the kingdom Plantae, *C. albicans, Cladosporium,* and *Saccharomyces* (**Fig. 2C**). These findings suggest that the gut mycobiota does not generate such a strong immune response compared to other bacterial and viral infections, *e.g.*, CMV^15^, SARS-CoV-2^17^ and *B. burgdorferi^16^*.

**Figure 2.**
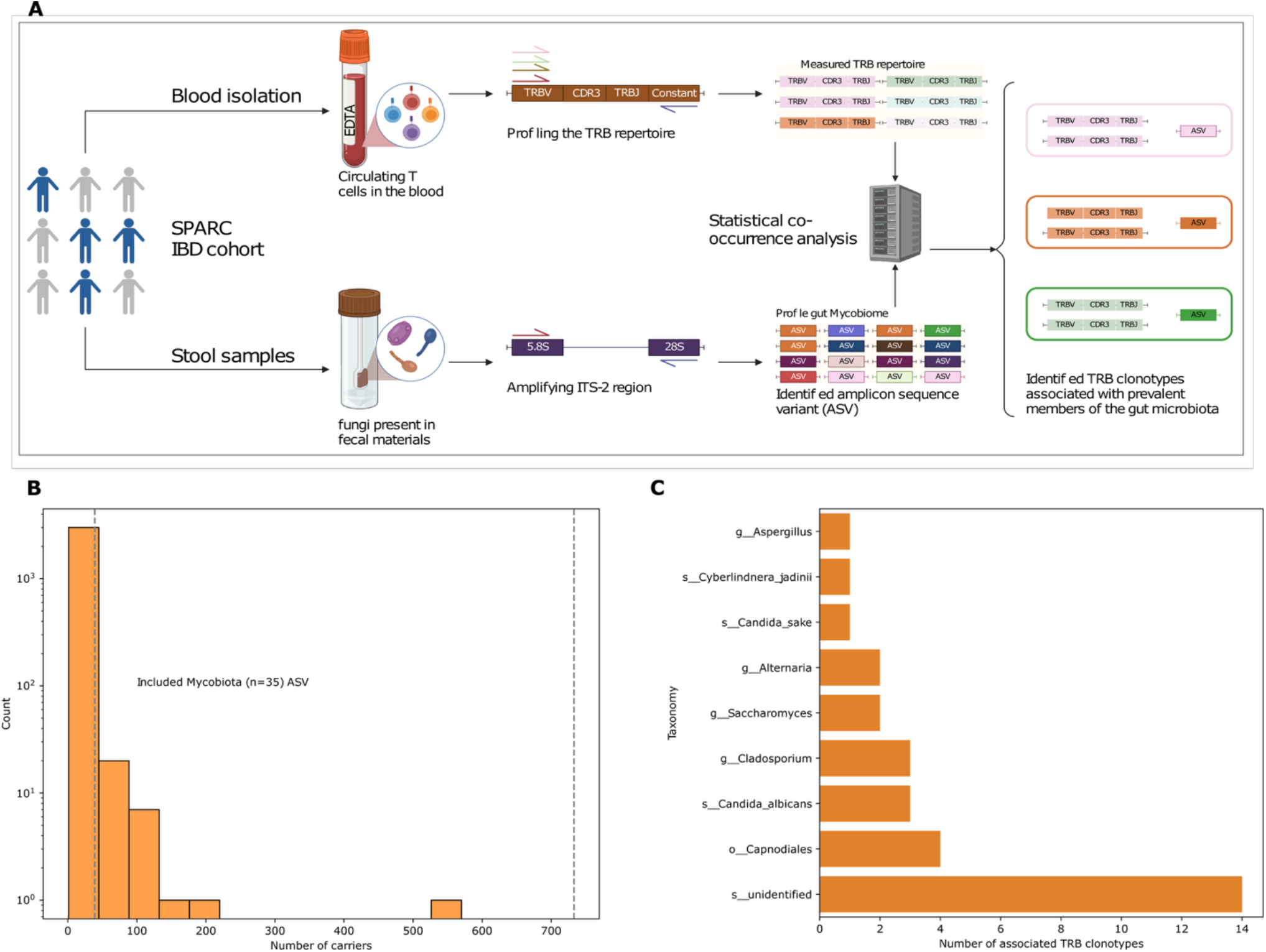
Mapping the interaction between the T cell repertoire and the gut mycobiome statistically. **(A)** shows the approach used for integrating the T cell repertoire dataset statistically with the ITS-2 to identify clonotypes associated with the presence of diMerent members of the gut mycobiota. (**B)** illustrates the private nature of the gut mycobiome, where most fungi are detected in only one individual, and only 35 ASVs were detected in more than 5% of the study cohort. (**C**) the number of clonotypes associated with diMerent members of the gut mycobiota, with the unidentified sequences representing diMerent members of plants identified via a BLAST search^49^, potentially derived from food.

### Mapping antibody responses toward *S. cerevisiae* in healthy controls and individuals with CD using PhIP-Seq

While a significant increase in anti-fungal clonotypes was detected in the T cell repertoire of individuals with CD, functional antibody repertoires showed elevated responses toward herpesviruses^29^ and bacterial flagellins in individuals with CD^26,30^. As this difference in detected responses can be attributed to a lack of representation of fungal antigens in the PhIP-seq library, we designed a novel phage library covering >12,000 peptides derived from 774 *S. cerevisiae’s* proteins (**Methods**). These proteins were mainly annotated to be either a membrane protein or secreted proteins^31,32^ as these sites might be a more plausible target for antibodies. Thus, we aimed with this library to narrow down the candidate peptides/antigens of *S. cerevisiae* as they remain unknown for CD. We also included the full proteome of other common viral pathogens such as Epstein Barr virus (EBV), and cytomegalovirus (CMV) as well as common vaccines such as the poliovirus vaccine, to quantify the overall immunogenetic properties of *S. cerevisiae* relative to these antigenic exposures (**Fig. S5**). The new PhIP-Seq library was then combined with the previous library^14,33^ and was then used to screen the repertoire of 100 individuals with CD and 60 healthy individuals (**Fig. 3A**).

**Figure 3:**
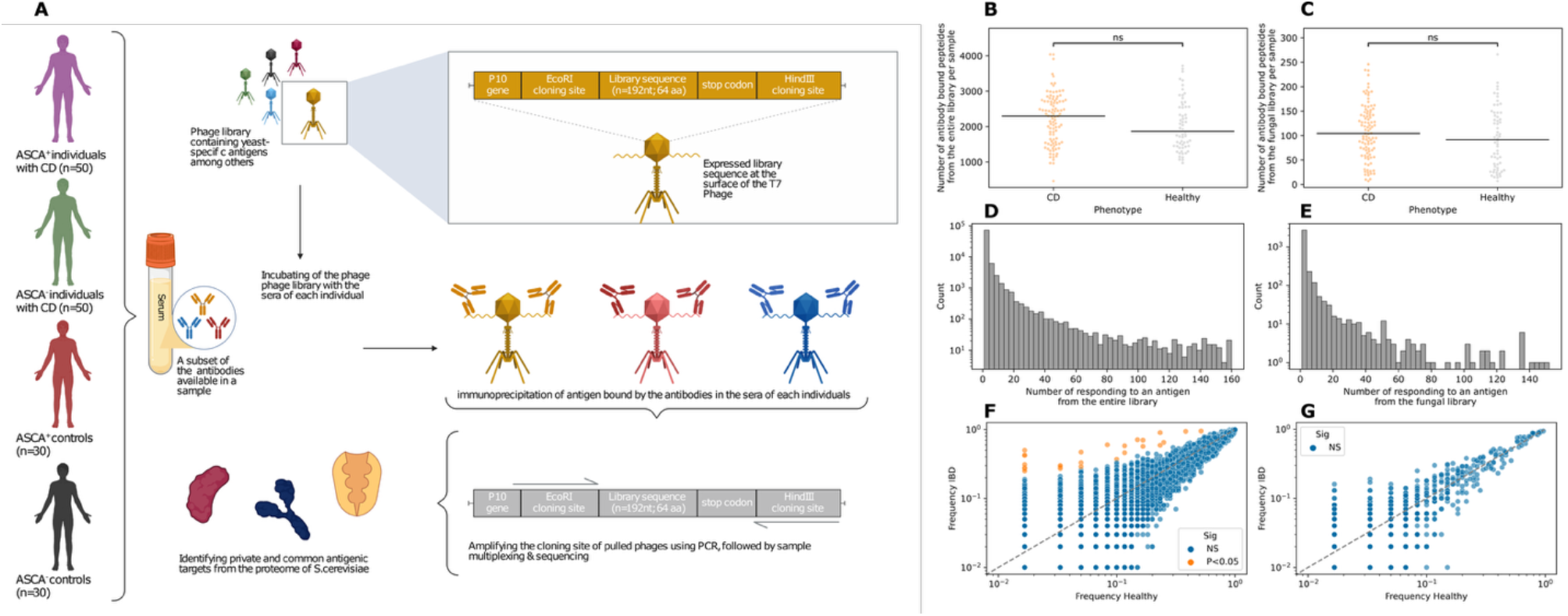
Identifying anti-fungal responses at the antibody levels using PhIP-Seq. (**A**) depicts the study design for identifying anti-fungal responses using PhIP-Seq by screening the sera of 100 individuals with CD and 60 healthy controls against a novel yeast library in combination with other existing libraries. (**B**) and (**C**) show the number of antigens per individual from the combined antigen libraries used in the study (**B**) or only the yeast-library (**C**). (**D**) and (**E**) shows the number of individuals binding a particular antigen from the entire library (**D**) or the fungal library (**E**). (**F**) shows antigenic exposures associated with CD predominantly, anti-flagellin responses, while (**F**) shows the lack of association between anti-fungal responses and CD as measured by PhIP-Seq. In both (**F**) and (**G**), the x-axis depicts the frequency of an antigen in healthy individuals, while the y-axis shows the frequency in individuals with CD. Also, orange dots represent antigens that have a higher prevalence in CD than expected by chance, as measured by Fisher’s exact test. In panels (**B**) and (**C**), a two-sided Mann-Whitney Wilcoxon U test was used to compare between the number of bound antigens between healthy controls and individuals with CD.

Similar to previous findings, there was not any significant difference in the number of bound antigens after including the fungal library between individuals with and without CD (**Fig. 3B-3C**). Focusing on the fungal antigens, most responses were private and were not shared among individuals with and without CD (**Fig. 3D-3E**). By comparing the functional antibody repertoire against the entire library, we detected elevated antibody responses toward flagellin antigens in individuals with CD relative to individuals without CD (**Fig. 3F**). Nonetheless, by focusing only on the fungal antigens, we did not detect any significant association (**Fig. 3G**). These findings indicate that anti-fungal responses represent a smaller subset of dysregulated immune responses in individuals with CD relative to the predominant anti-flagellin signal.

To follow this further, we focused on the yeast library and compared the prevalence of anti- *S. cerevisiae* antigens relative to other known viral pathogens, bacterial virulence factors and common vaccines (**Fig. 4A**). In comparison to prevalent infections such as EBV and CMV, the prevalence of anti- *S. cerevisiae* responses was significantly lower, P= 3.3×10^−20^, and 1.2×10^−11^, respectively. Furthermore, it was also much lower than the virulence factors derived from common bacterial pathogens, namely, *K. pneumonia* and *V. cholera* (P= 7.6×10^−5^). Similarly, it also had a much lower prevalence relative to common vaccines, *e.g.* Poliovirus (P= 6.8×10^−4^). Indeed, anti- *S. cerevisiae* responses had a comparable prevalence to that of negative controls such as the Cyanophage or even random sequences.

**Figure 4:**
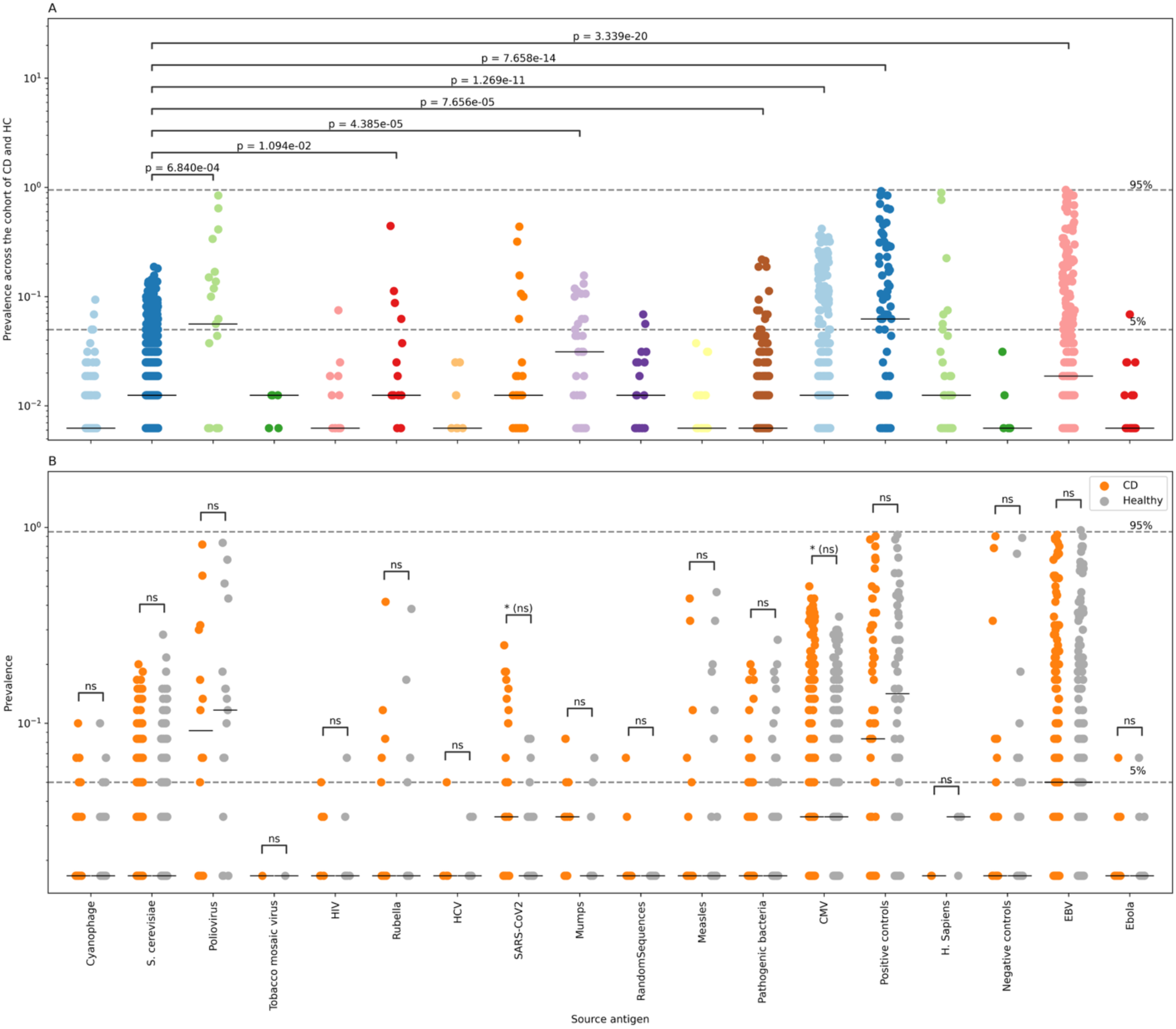
Characterization of the immunogenic properties of S. cerevisiae relative to other antigenic sources and among individuals with and without CD. (**A**) the prevalence of immune responses directed against diMerent pathogens included in the novel phage library reported in the current study. (**B**) a comparison between the prevalence of immune responses toward the diMerent antigenic sources included in the library in healthy individuals and individuals with CD, with only statistically significant diMerences shown. In both panels, the Mann-Whitney U test was used to compare the diMerent groups, and the Benjamini-Hochberg correction was used to correct for multiple testing.

Out of 1,956 peptides from the proteome of *S. cerevisiae* that were able to elicit an immune response in at least one individual, only 83 peptides generated an immune response that was detected in >5% of all study participants, *i.e.* eight donors. The most antigenic peptide was derived from the PSH1 protein^34^, *generating* the most prevalent response in ∼19% of individuals, followed by an antigenic peptide from the PAN1 proteins^35^ at ∼18% and lastly from the LSB6 proteins^36^ at ∼15%. These results corroborated previous findings that most *S. cerevisiae* proteins fail to mount a strong immune response across multiple individuals as seen with other viral and pathogenic bacteria. Lastly, we aimed at comparing the prevalence of responses toward these antigens in individuals with CD relative to healthy controls. Thus, we down sampled the number of individuals with CD to the same number of healthy controls, *i.e.* 60 individuals and subsequently compared the prevalence in response between these two groups (**Fig. 4B**). Across all antigenic sources included in the library, we observed a compared level of prevalence between individuals with CD and healthy controls. Suggesting the variation in the immunogenic properties of an exposure is greater than the difference in the mounted response between individuals with CD and healthy controls.

### The repertoire of ASCA^+^ individuals harbors multiple alterations relative to that of ASCA^-^ individuals

We then aimed to investigate the relationship between ASCA-positivity and its impact on the antibody repertoire. Given that our study contains 80 ASCA^+^ individuals (50 individuals with CD and 30 healthy controls) as well as 80 ASCA^-^ individuals (50 individuals with CD and 30 healthy controls), we compared the binding to the different pathogens included in the fungal-focused PhIP-Seq library. Focusing only on individuals with CD, we observed a comparable level of immune responses toward the different organisms included in the library (**Fig. S6A**). Then we extracted only antigens derived from *S. cerevisiae* and we compared the frequency of immune responses targeting them in ASCA^+^ and ASCA^-^ individuals with CD (**Fig. S6B**), as well as in ASCA^+^ and ASCA^-^ controls (**Fig. S6C**). In addition, to all ASCA^+^ and ASCA^-^ individuals regardless of their disease status (**Fig. S6D**). Across these the groups, we could not detect any peptide antigen from *S. cerevisiae* that was significantly associated with ASCA-positivity.

Then we extended this analysis to the entire antigenic library, included the antigens reported by Vogl *et al.^14^* and Leviatan *et al.^33^*. By comparing the repertoires of ASCA^+^ and ASCA^-^ individuals with CD we identified 44 antigenic peptides that were significantly prevalent in ASCA^+^ individuals relative to ASCA^-^ individuals (**Fig. 5A**) and 17 antigenic exposures that were significantly more prevalent in ASCA^-^ individuals relative to ASCA^+^ individuals (**Fig. 5A**). However, by comparing the repertoire of ASCA^+^ and ASCA^-^ controls we could not detect any antigenic peptide that were associated with the ASCA status, potentially due to the small sample size of healthy controls (**Fig. 5B**). By comparing the repertoire of all ASCA^+^ individuals to that of ASCA^-^ individuals, we detected 64 antigens that were associated with ASCA-positivity (**Fig. 5C**) and 30 antigens that were enriched in ASCA- individuals (**Fig. 5C**).

**Figure 5:**
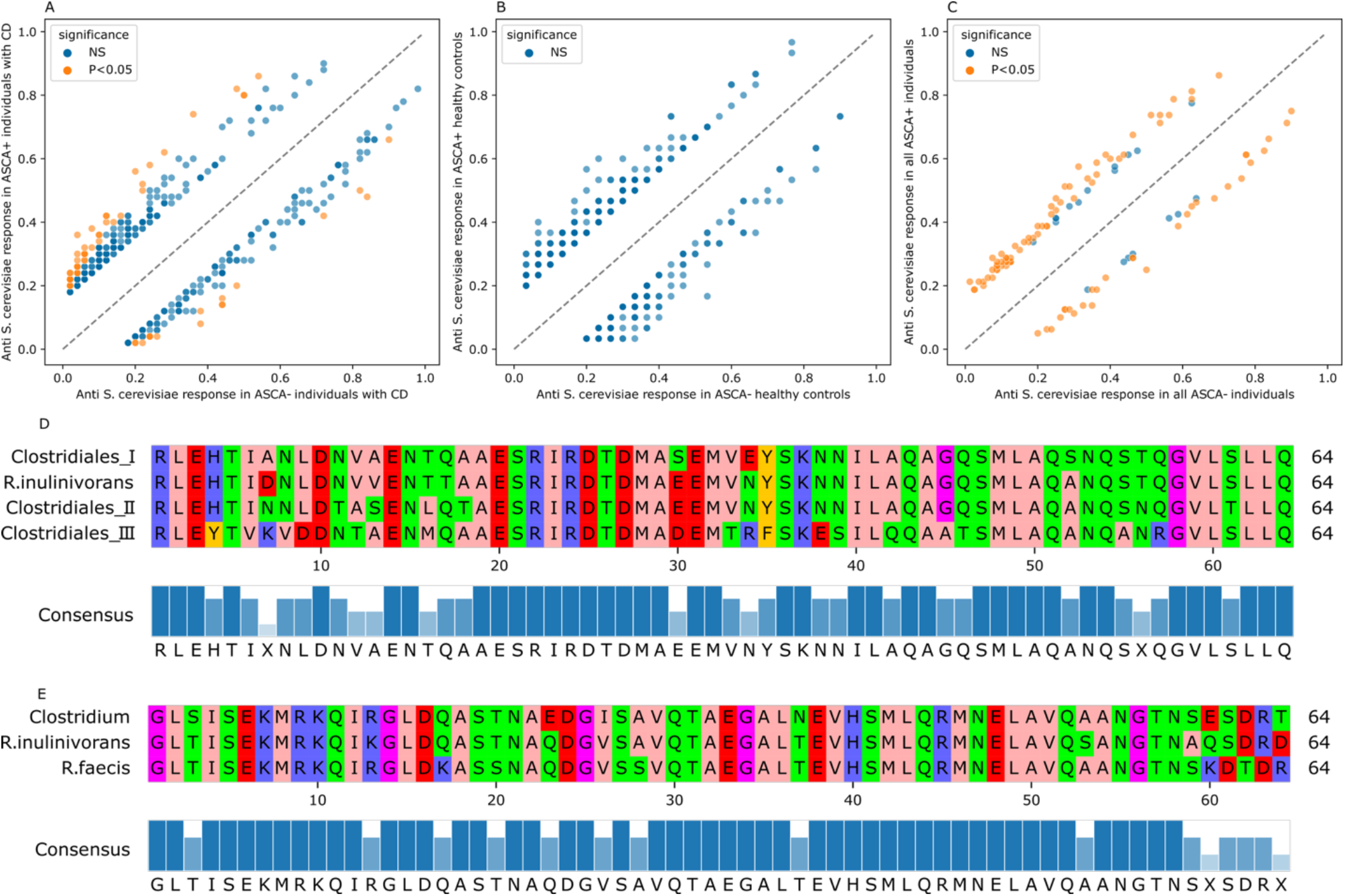
antigenic exposures associated with ASCA-positivity. (**A**) a comparison between the immune repertoire of ASCA^+^ and ASCA^-^ individuals with CD, similarly (**B**) depicts the same relationship in ASCA^+^ and ASCA^-^ controls. Lastly, (**C**) depicts the diMerence in antigenic binding between all ASCA^+^ and ASCA^-^ individuals regardless of disease status. (**D**) and (**E**) represent the sequence motif of the two main flagellin antigens associated with ASCA-positivity. In (**A**-**C**), we selected antigens detected in both ASCA^+^ and ASCA^-^ individuals and with a diMerence in prevalence between them of more than 15%. After that we used the Fisher’s exact test to compare their prevalence in cases and controls, lastly, we used the Benjamini-Hochberg approach to correct for multiple testing.

To investigate antigens associated with ASCA-positivity, we focused on the union of all antigens that were associated with ASCA-positivity in the analyses above (**Fig. 5A**-**5C**). One of the dominant signals was derived from bacterial flagellin predominantly from several members of the Clostridiales class and particularly from the Lachnospiraceae family, such as *R. inulinivorans* and *R. faecis.* These anti-flagellin responses were derived from two main distinct sequence clusters (**Fig. 5D-5E**), thus, suggesting that the flagella of these microbiota are a major antigenic target in ASCA^+^ individuals relative to ASCA^-^ individuals. Beside the increase in anti-flagellin response, we also observed an increase in autoantibodies in ASCA^+^ individuals. These antibodies were targeting several host proteins such as Ryanodine receptor 3, Thyroglobulin and large aggregating cartilage proteoglycan core protein. Lastly, we observed an increase in antibody responses against several bacteriophages such as the Enterobacter phage, the Lactobacillus phage, and the Klebsiella phage. There was a fewer number of antigens that were enriched in ASCA^-^ individuals, these antigens were mainly derived from different bacterial virulence factors mainly from *S. pneumoniae, S. pyogenes* and pathogenic *E. coli* strains.

## Discussion

While elevated T cell responses toward fungal antigens have been previously identified in individuals with CD^8^, a quantification of the degree of expansion at the population scale and relative to other known antigenic exposures, such as flagellins, has not been performed. Thus, we started the analysis by studying the expansion of these clonotypes in a large cohort of individuals containing 2,804 individuals with IBD from the SPARC IBD cohort. This enabled us to identify a significant expansion of anti-fungal clonotypes in individuals with CD relative to UC, specifically, toward *S. cerevisiae* and particularly in individuals with a penetrating disease. Nonetheless, the expansion of these clonotypes varied significantly, *i.e.,* by two orders of magnitude, within individuals with CD. While this might indicate heterogeneity in the burden of anti-fungal responses in individuals with and without CD specifically as the ASCA status of these individuals is not known, it can also be attributed to the small sample size used to identify anti- fungal responses in the Martini *et al.*^8^ study (n=6 individuals). Thus, a small subset of anti-fungal clonotypes that are restricted to specific human leukocyte antigens (HLA) alleles, ^8^could be identified. Hence, the apparent heterogeneity in response might reflect the similarity in the HLA background of individuals from the SPARC IBD cohort and the Martini *et al.* study cohort^8^.

We also aimed to statistically link TCR clonotypes with the gut mycobiome by analyzing co-occurrence patterns between T cell repertoires and the gut mycobiota profiled using ITS2. However, a major challenge was the low prevalence of most gut mycobiota, where only 35 ASVs were detected in >5% of the study cohort. Furthermore, most of the ASVs were likely derived from dietary intakes, such as vegetables or food processing fungi, such as *S. cerevisiae* and different *Candida* species^37^. These findings indicate that most of these mycobiota are transitioning through the gut but are rarely colonizing the gut, with a minimal interaction with the host immune system. Indeed, there was no correlation between the levels of ASCA antibodies, which targets fungal mannoproteins, and the presence of *S. cerevisiae* at colonic mucosa^38^. Furthermore, the levels of ASCA appears to be stable across different disease trajectories, as well as therapeutic and surgical interventions^39^. These observations indicate a decoupling between the stable immune repertoire and the more dynamic gut mycobiome which heavily influenced by food intake^40^, hence, linking the prevalence of these mycobiota with a system that has a long-lived memory, such as the immune repertoire, is challenging. Nonetheless, given the less diverse gut mycobiome, functional antigen-stimulation assays and TCR-Seq^41,42^ can be used to study or identify the anti- fungal compartment of the T cell repertoire.

The antigenicity of these mycobiota, *e.g. S. cerevisiae,* appears to be much lower than other prevalent viral infections such as EBV, CMV or even common vaccines such as the polio vaccine as observed in our PhIP-Seq experiment. While we did not include all *S. cerevisiae* proteins in the current library, we included a representative set of mainly secreted and membrane proteins (n=774 proteins). As PhIP-Seq can only be used to identify linear antigens without post-translational modifications (PTMs) recognized by the sera’s antibodies, confirmational antigens with or without PTMs cannot be identified using PhIP-Seq^22^. While this might explain the low prevalent immune responses toward fungal antigens, other pathogens included in the library such as EBV which encodes for multiple glycoproteins, have generated a robust and prevalent immune response as measured by the PhIP-Seq.

A striking observation was the lack of any association between anti- S. *cerevisiae* responses and ASCA status which showed elevated immune responses toward other antigens such as bacterial flagellins from the Clostridales order. ASCA-positivity correlates with many important clinical features of CD such as, earlier disease-onset, ileal involvement and an overall more aggressive disease course^39^. Hence, these ASCA-positive associated antigenic exposures might rather be associated with a specific clinical feature such as disease location, onset or severity that is more common in ASCA^+^ individuals relative to ASCA^-^ individuals. Higher ASCA levels was also observed in other inflammatory diseases of the small intestine such as celiac disease^43,44^ where gluten has been shown to be the main driver of the disease. While the cause of ASCA-positivity remains to be identified, but potential a more defective gut barrier, as seen in CD or celiac disease, might allow the diffusion of some fungal polysaccharides into submucosal tissues where it might derive a strong antibody response toward this immunogenetic polysaccharide^45^.

The presence of ASCA antibodies strongly correlates with several other serological markers^46^, such as CBir1^47^, Fla2 and FlaX^48^ which are derived from different flagellin proteins, corroborating our findings about elevated anti-flagellin responses in ASCA^+^ individuals with CD. These surface proteins such as the flagellins and mannoproteins, might be detaching from their source organism, diffuse deeper into submucosal tissues because of a defective barrier to elicit a strong immune response without being actively implicated in the disease. Hence, elevated anti-fungal responses might be a marker of a more aggressive disease in the small intestine, *i.e.* a consequence of the disease, but it does not imply a causal role of these fungal exposures in the disease.

In conclusion, we aimed to perform an exhaustive investigation of immune responses toward the gut mycobiome in individuals with IBD. While our results point toward an increase in anti-fungal T cell responses in individuals with IBD, the magnitude of this expansion is significantly smaller than other antigenic exposures, such as bacterial flagellins which are significantly and more robustly associated with CD^25^.

## Methods

### Cohort description and ethical approval

Multiple cohorts were utilized in the current study, primarily the SPARC IBD cohort^27^, which is maintained by the IBD Plexus research program of the Crohn’s & Colitis Foundation.

### Profiling and processing the TRB repertoire

The TRB repertoire of the SPARC IBD cohort was profiled using up to 18ug of DNA isolated from peripheral blood through the immuneSEQ assay (Adaptive Biotechnologies)^17,50^. After clonotype identification, non-productive clonotypes, *i.e.,* clonotypes containing a stop codon or frameshift and thus cannot produce a functional TRB chain, were filtered. After that, we collapsed the TRB chain with the same V and J genes and the complementarity-determining region 3 (CDR3) amino acid sequence together and summed their different expansion level.

### Profiling the gut mycobiome using ITS2

All methods related with ITS2 sequencing analysis were retrieved from Diversigen^SM^. Genomic DNA extraction was performed using the Qiagen Mag Attract Power Soil kit (cat. No.: 27000-4- EP) following the manufacturer’s instructions. The ITS2 region was amplified using Invitrogen AccuPrime High Fidelity kit (cat. No.: 12346094). PCR mix contain 10.05 μL water, 2 μL 10X reaction buffer, 0.15 μL Accuprime Taq, 0.8 μL BSA, 5 μL template DNA, 2 μL primer mix (4 μM). Target-specific primer sequences are provided for each PCR primer: 5’- GCATCGATGAAGAACGCAGC-3’ (ITS3F), 5’-TCCTCCGCTTATTGATATGC-3’ (ITS4). The primers also contain adapter sequences and single-end barcodes (not provided by Diversigen^SM^). PCR starts from initial denaturation at 95°C for 2 min followed by 35 amplification cycles of 20 s at 95°C, 45 s at 56°C, and 90 s at 72°C followed by final extension at 72°C for 10 min. The purification of PCR product is conducted using the QIAquick PCR purification kit (cat. No.: 28104) and the yield is quantified using Invitrogen Quant-iT Picogreen dsDNA assay kit (cat. No.: P7589). Amplicons are pooled and cleaned using the Invitrogen charge switch kit. Sequencing was performed in the Illumina MiSeq platform using the 2×300 bp paired-end protocol.

The read pairs are demultiplexed based on their unique molecular barcodes, denoised and merged using DADA2^51^ and chimera removal is conducted using VSEARCH^52^. ITS2 sequences are clustered into Operational Taxonomic Units (OTUs) at a similarity cutoff value of 97%. Taxonomy assignment was performed using the scikit-learn classifier and optimized variable region-specific version of UNITE Database^53^. After taxonomy assignment, a rarefied OTU table is generated from the output files in the previous two steps for downstream analysis.

### Discovering mycobiota-associated TRB clonotypes

We focused this analysis on a subset of individuals from the SPARC IBD cohort with paired TRB and ITS2 measurements (n=637). We started by identifying public members of the gut mycobiome, *i.e.,* mycobiota with a prevalence between 5 and 95% of the cohort. Given the private and dynamic nature, only 35 amplicon sequence variants (ASVs) fall within the defined prevalence range. For each ASV, we split the cohort into carriers and non-carriers and then used a one-sided Fisher’s exact test to identify public clonotypes associated with the carriership of the identified ASV^15^. Lastly, we used a Bonferroni-based correction method to identify correct for multiple-testing where a clonotype was linked to a particular gut mycobiota if the association P- value was less than 2.8×10^−6^ which corresponds to the wildly used P-value of (1×10^−4^)^16,17,54^ divided by the number of ASV-tested (n=35).

### Designing a novel library covering the proteome of *S. cerevisiae*

To build a library of potential antigens from *S. cerevisiae,* we downloaded the pan proteome of *S. cerevisiae* from UniProt^31^ and selected secreted and surface membrane proteins. Subsequently, we performed *in silico* digestion using a sliding window approach of length 64 amino acids and an overlap of 20 amino acids. We also added known positive controls, such as the proteome of common viral pathogens, such as Mumps, Measles, Rubella, Epstein-Barr virus, and cytomegalovirus. We also included multiple negative controls, such as human proteins and non- human pathogens, and previously utilized negative controls^14^. The proteome of these positive and negative controls were combined with the pan proteome of *S. cerevisiae* and *in silico* digested using the same approach. To remove redundancies in the generated peptides, clustering using cd-hit^55^ was performed, and subsequently, sequences were reverse-translated *in silico* into DNA. During reverse translation, the codon usage was optimized for protein-translation in *E. coli,* and any EcoRI and Hind-III restriction site was removed. After that, the following adaptor sequences were added as a prefix (*GATGCGCCGTGGGAATTCT*) and a suffix (*TGAAAGCTTGCCACCCGAC*) to the reverse-translated sequences^14^, generating a 230nt DNA sequence. Lastly, the generated 15,000 oligos were commercially synthesized using Agilent’s SureDesign platform.

### Cloning the *S. cerevisiae* library and phage-immunoprecipitation sequencing

After ordering the library, it was amplified using PCR in accordance with the manufacturer’s instructions. Subsequently, 1 μg of the amplified library was incubated with the *EcoRI and HindIII* (FastDigest; Thermo Fisher) according to the manufacturer’s instructions. After that, the generated product was cleaned using magnetic beads (AMPure XP Beads; Beckman Coulter) to remove DNA fragments shorter than 150bp, salts, and deactivated enzymes. The cleaned product (202bp with four sticky-ends from each end) was then cloned into the T7 Select 10-3b vector (Merck Millipore, T7Select^®^10-3 cloning kit, product no. 70550) according to the manufacturer’s instructions. The cloning reaction was stopped after 16 hrs, and *in vitro* packaging (Merck Millipore, T7Select^®^ Packaging Kit, product no. 70014) was used to assemble the cloned production into phages. These T7 phages were then amplified using a liquid-media amplification assay in accordance with the manufacturer’s instructions (Merck Millipore), followed by a titration assay to estimate the yield of the amplified phages. After that, one-stock of phages was amplified using a plate-based amplification approach according to the manufacturer’s instructions (Merck Millipore) to amplify the phages. Lastly, the produced library was then combined with previously described libraries^14,33^ to generate the input phage library, and phage-immunoprecipitation sequencing was conducted on this joint library as discussed previously^14^.

### Identifying antibody-bound fungal antigens

After sequencing and demultiplexing, the generated reads were down sampled into 1,250,000 reads using seqtk. Subsequently, the reads were aligned to the reference library using bowtie (v 1.0)^56^. Then we considered paired reads that aligned perfectly to only one oligo without any mismatches to generate the count matrix, which contains the number of reads aligned to each oligo in each sample. Subsequently, we modeled the generated counts using a generalized Poisson model parametrized by a rate λ and a dispersion factor θ. These parameters were estimated from the counts of the generated phages, *i.e.,* input samples, and immunoprecipitation of the same input library without any antibody enrichment, *i.e.,* mocked samples^14^. After estimating λ and θ for the expected number of reads from an immunoprecipitation pulldown of the input library, *i.e.,* the null distribution, a generalized Poisson model was used to estimate the probability of observing a particular count for a given oligo in a sample under the same null distribution. Hence, a smaller P-value indicates that observed counts are less likely under the null distribution, and hence the count was higher due to antibody enrichment. We used a Bonferroni-based correction (α = 0.05) to identify the bound antigen in each sample.

## Acknowledgments

We would also like to thank the Crohn’s & Colitis Foundation IBD Plexus program for providing us with the T cell repertoire profiles and the genotypes of the SPARC IBD cohort. The project was funded by the EU Horizon Europe Program grant *miGut-Health: personalized blueprint of intestinal health* (101095470) and the EU program for Research and Innovation “Horizon Health” (HORIZON-HLTH-2023-DISEASE-03) *ID-DarkMatter-NCD* (897856542). Additionally, the project received funding from the German Research Foundation (DFG) Research Unit 5042: miTarget – The Microbiome as a Therapeutic Target in Inflammatory Bowel Diseases along with funding from the DFG Cluster of Excellence 2167 “Precision Medicine in Chronic Inflammation (PMI)”. The SPARC IBD cohort is maintained by the Crohn’s & Colitis Foundation for research use.

## Reference

1. Hampe, J. et al. A genome-wide association scan of nonsynonymous SNPs identifies a susceptibility variant for Crohn disease in ATG16L1. Nat Genet 39, 207–211 (2007).

2. Ogura, Y. et al. A frameshift mutation in NOD2 associated with susceptibility to Crohn’s disease. Nature 411, 603–606 (2001).

3. Hugot, J.-P. et al. Association of NOD2 leucine-rich repeat variants with susceptibility to Crohn’s disease. Nature 411, 599–603 (2001).

4. Goyette, P. et al. High-density mapping of the MHC identifies a shared role for HLA-DRB1*01:03 in inflammatory bowel diseases and heterozygous advantage in ulcerative colitis. Nat Genet 47, 172–179 (2015).

5. Degenhardt, F. et al. Trans-ethnic analysis of the human leukocyte antigen region for ulcerative colitis reveals shared but also ethnicity-specific disease associations. medRxiv 2020.07.29.20162552 (2020) doi:10.1101/2020.07.29.20162552.

6. Ebert, A. C. et al. Risk of inflammatory bowel disease following hospitalisation with infectious mononucleosis: nationwide cohort study from Denmark. Nat Commun 15, 8383 (2024).

7. Uchida, A. M. et al. Escherichia coli–Specific CD4+ T Cells Have Public T-Cell Receptors and Low Interleukin 10 Production in Crohn’s Disease. Cell Mol Gastroenterol Hepatol 10, 507–526 (2020).

8. Martini, G. R. et al. Selection of cross-reactive T cells by commensal and food-derived yeasts drives cytotoxic TH1 cell responses in Crohn’s disease. Nat Med 29, 2602–2614 (2023).

9. Sendid, B. et al. Specific antibody response to oligomannosidic epitopes in Crohn’s disease. Clinical Diagnostic Laboratory Immunology 3, 219–226 (1996).

10. Sun, M., Ju, J., Xu, H. & Wang, Y. Intestinal fungi and antifungal secretory immunoglobulin A in Crohn’s disease. Front Immunol 14, (2023).

11. Nelson, A. et al. The Impact of NOD2 Genetic Variants on the Gut Mycobiota in Crohn’s Disease Patients in Remission and in Individuals Without Gastrointestinal Inflammation. J Crohns Colitis 15, 800–812 (2021).

12. Ott, S. J. et al. Fungi and inflammatory bowel diseases: Alterations of composition and diversity. Scand J Gastroenterol 43, 831–841 (2008).

13. Mahdy, A. K. H. et al. Bulk T cell repertoire sequencing (TCR-Seq) is a powerful technology for understanding inflammation-mediated diseases. J Autoimmun 149, 103337 (2024).

14. Vogl, T. et al. Population-wide diversity and stability of serum antibody epitope repertoires against human microbiota. Nat Med 27, 1442–1450 (2021).

15. Emerson, R. O. et al. Immunosequencing identifies signatures of cytomegalovirus exposure history and HLA-mediated ekects on the T cell repertoire. Nat Genet 49, 659–665 (2017).

16. Greissl, J. et al. Immunosequencing of the T-Cell Receptor Repertoire Reveals Signatures Specific for Identification and Characterization of Early Lyme Disease. medRxiv 2021.07.30.21261353 (2022) doi:10.1101/2021.07.30.21261353.

17. Gittelman, R. M. et al. Longitudinal analysis of T cell receptor repertoires reveals shared patterns of antigen-specific response to SARS-CoV-2 infection. JCI Insight 7, (2022).

18. Pesesky, M. et al. Antigen-driven expansion of public clonal T cell populations in inflammatory bowel diseases. J Crohns Colitis jjaf048 (2025) doi:10.1093/ecco-jcc/jjaf048.

19. Elabd, H., Mahdy, A. & Franke, A. OP11 Analysing the T cell receptor beta chain repertoire of 2,800 Inflammatory Bowel Disease patients identifies public T cell responses involved in the pathogenesis of Crohn’s Disease and Ulcerative Colitis and quantifies the impact of surgery and therapy on the immune repertoire of IBD patients. J Crohns Colitis 19, i21–i23 (2025).

20. Mahdy, A. et al. P0125 Crohn’s-associated invariant T Cells are associated with disease severity and location and are not akected by medication intake. J Crohns Colitis 19, i507–i508 (2025).

21. Rawat, P. et al. Identification of a type 1 diabetes-associated T cell receptor repertoire signature from the human peripheral blood. medRxiv 2024.12.10.24318751 (2024) doi:10.1101/2024.12.10.24318751.

22. Mohan, D. et al. PhIP-Seq characterization of serum antibodies using oligonucleotide-encoded peptidomes. Nat Protoc 13, 1958–1978 (2018).

23. Andreu-Sánchez, S. et al. Phage display sequencing reveals that genetic, environmental, and intrinsic factors influence variation of human antibody epitope repertoire. Immunity 56, 1376–1392.e8 (2023).

24. Klompus, S. et al. Cross-reactive antibodies against human coronaviruses and the animal coronavirome suggest diagnostics for future zoonotic spillovers. Sci Immunol 6, eabe9950 (2021).

25. Bourgonje, A. R., Hörstke, N. V, Fehringer, M., Innocenti, G. & Vogl, T. Systemic antibody responses against gut microbiota flagellins implicate shared and divergent immune reactivity in Crohn’s disease and chronic fatigue syndrome. Microbiome 12, 141 (2024).

26. Bourgonje, A. R. et al. Phage-display immunoprecipitation sequencing of the antibody epitope repertoire in inflammatory bowel disease reveals distinct antibody signatures. Immunity 56, 1393–1409.e6 (2023).

27. Rakals, L. E. et al. The Development and Initial Findings of A Study of a Prospective Adult Research Cohort with Inflammatory Bowel Disease (SPARC IBD). Inflamm Bowel Dis 28, 192–199 (2022).

28. ElAbd, H. et al. Decoding the restriction of T cell receptors to human leukocyte antigen alleles using statistical learning. bioRxiv 2022–2025 (2025).

29. Nandy, A. et al. Epstein-Barr Virus (EBV) Exposure Precedes Crohns Disease Development. Gastroenterology (2025) doi:10.1053/j.gastro.2025.01.247.

30. Lodes, M. J. et al. Bacterial flagellin is a dominant antigen in Crohn disease. J Clin Invest 113, 1296–1306 (2004).

31. Consortium, T. U. UniProt: the Universal Protein Knowledgebase in 2025. Nucleic Acids Res 53, D609–D617 (2025).

32. Bateman, A. UniProt: A worldwide hub of protein knowledge. Nucleic Acids Res (2019) doi:10.1093/nar/gky1049.

33. Leviatan, S. et al. Allergenic food protein consumption is associated with systemic IgG antibody responses in non-allergic individuals. Immunity 55, 2454–2469.e6 (2022).

34. Hewawasam, G. et al. Psh1 Is an E3 Ubiquitin Ligase that Targets the Centromeric Histone Variant Cse4. Mol Cell 40, 444–454 (2010).

35. Kamińska, J., Wysocka-Kapcińska, M., Smaczyńska-de Rooij, I., Rytka, J. & Żołądek, T. Pan1p, an actin cytoskeleton-associated protein, is required for growth of yeast on oleate medium. Exp Cell Res 310, 482–492 (2005).

36. Han, G.-S., Audhya, A., Markley, D. J., Emr, S. D. & Carman, G. M. The Saccharomyces cerevisiae LSB6 Gene Encodes Phosphatidylinositol 4-Kinase Activity*. Journal of Biological Chemistry 277, 47709–47718 (2002).

37. Kieliszek, M. et al. Biotechnological use of Candida yeasts in the food industry: A review. Fungal Biol Rev 31, 185–198 (2017).

38. Mallant-Hent, R. C. et al. Correlation between Saccharomyces cerevisiae DNA in intestinal mucosal samples and anti-Saccharomyces cerevisiae antibodies in serum of patients with IBD. World journal of gastroenterology: WJG 12, 292 (2006).

39. Sendid, B. et al. From ASCA breakthrough in Crohn’s disease and Candida albicans research to thirty years of investigations about their meaning in human health. Autoimmun Rev 23, 103486 (2024).

40. A, A. T. et al. Investigating Colonization of the Healthy Adult Gastrointestinal Tract by Fungi. mSphere 3, 10.1128/msphere.00092-18 (2018).

41. Bacher, P. et al. Antigen-reactive T cell enrichment for direct, high-resolution analysis of the human naive and memory Th cell repertoire. The Journal of Immunology 190, 3967–3976 (2013).

42. Klinger, M. et al. Multiplex Identification of Antigen-Specific T Cell Receptors Using a Combination of Immune Assays and Immune Receptor Sequencing. PLoS One 10, e0141561. (2015).

43. Granito, A. et al. Anti-Saccharomyces cerevisiae and perinuclear anti-neutrophil cytoplasmic antibodies in coeliac disease before and after gluten-free diet. Aliment Pharmacol Ther 21, 881–887 (2005).

44. Barta, Z., Csípõ, I., Szabó, G. G. & Szegedi, G. Seroreactivity against Saccharomyces cerevisiae in patients with Crohn’s disease and celiac disease. World J Gastroenterol 9, 2308 (2003).

45. Parikh, K. et al. Colonic epithelial cell diversity in health and inflammatory bowel disease. Nature 567, 49–55 (2019).

46. Gaifem, J. et al. A unique serum IgG glycosylation signature predicts development of Crohn’s disease and is associated with pathogenic antibodies to mannose glycan. Nat Immunol 25, 1692–1703 (2024).

47. Targan, S. R. et al. Antibodies to CBir1 Flagellin Define a Unique Response That Is Associated Independently With Complicated Crohn’s Disease. Gastroenterology 128, 2020–2028 (2005).

48. Schoepfer, A. M. et al. Phenotypic Associations of Crohn’s Disease with Antibodies to Flagellins A4-Fla2 and Fla-X, ASCA, p-ANCA, PAB, and NOD2 Mutations in a Swiss Cohort. Inflamm Bowel Dis 15, 1358–1367 (2009).

49. Altschul, S. F., Gish, W., Miller, W., Myers, E. W. & Lipman, D. J. Basic local alignment search tool. J Mol Biol 215, 403–410 (1990).

50. Carlson, C. S. et al. Using synthetic templates to design an unbiased multiplex PCR assay. Nat Commun 4, 2680 (2013).

51. Callahan, B. J. et al. DADA2: High-resolution sample inference from Illumina amplicon data. Nat Methods 13, 581–583 (2016).

52. Rognes, T., Flouri, T., Nichols, B., Quince, C. & Mahé, F. VSEARCH: a versatile open source tool for metagenomics. PeerJ 4, e2584 (2016).

53. Nilsson, R. H. et al. The UNITE database for molecular identification of fungi: handling dark taxa and parallel taxonomic classifications. Nucleic Acids Res 47, D259–D264 (2019).

54. Pesesky, M. et al. Antigen-driven expansion of public clonal T cell populations in inflammatory bowel diseases. bioRxiv 2024.05.15.594220 (2024) doi:10.1101/2024.05.15.594220.

55. Li, W. & Godzik, A. Cd-hit: a fast program for clustering and comparing large sets of protein or nucleotide sequences. Bioinformatics 22, 1658–1659 (2006).

56. Langmead, B., Trapnell, C., Pop, M. & Salzberg, S. L. Ultrafast and memory-efficient alignment of short DNA sequences to the human genome. Genome Biol 10, R25 (2009).

