## Supplementary figures for "Mapping adaptive immune responses toward fungal antigens in inflammatory bowel disease using T cell repertoire sequencing and phage-immunoprecipitation sequencing"

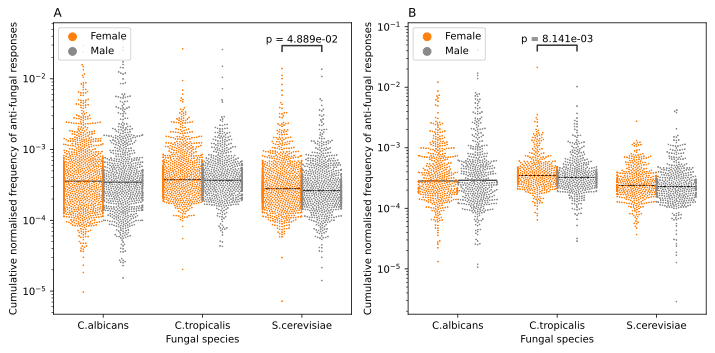

**Figure S1**: The interaction between biological sex and the expansion of fungal-specific clonotypes. (**A**) illustrate the expansion of anti- S. cerevisiae in Females with CD relative to males, while (**B**) shows the expansion of anti- C. tropicalis responses in females with ulcerative colitis relative to males.

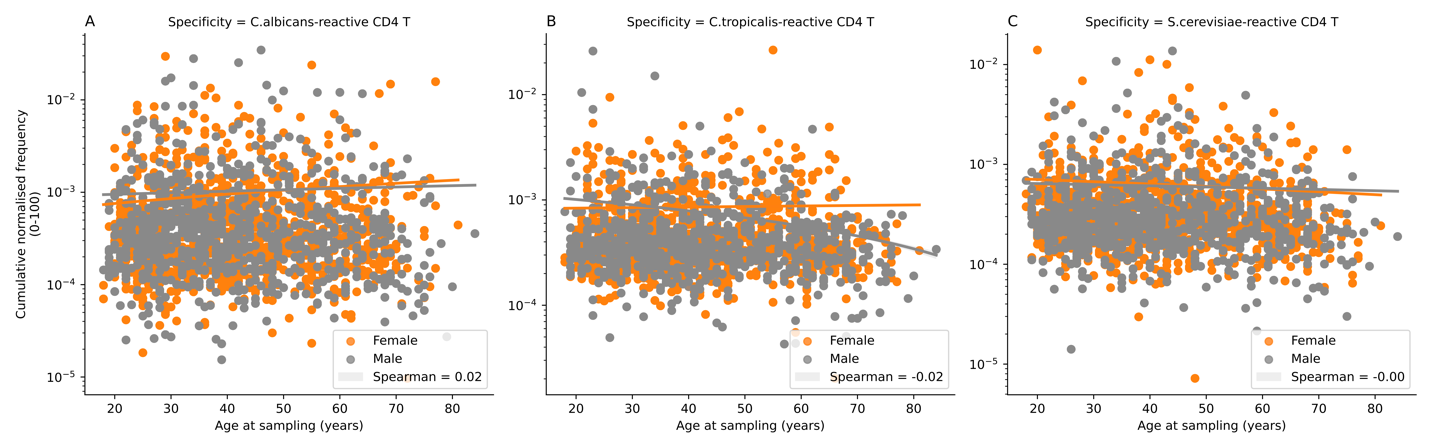

**Figure S2**: The relationship between the expansion of fungal-specific clonotypes and the biological age in individuals from the SPARC IBD cohort.

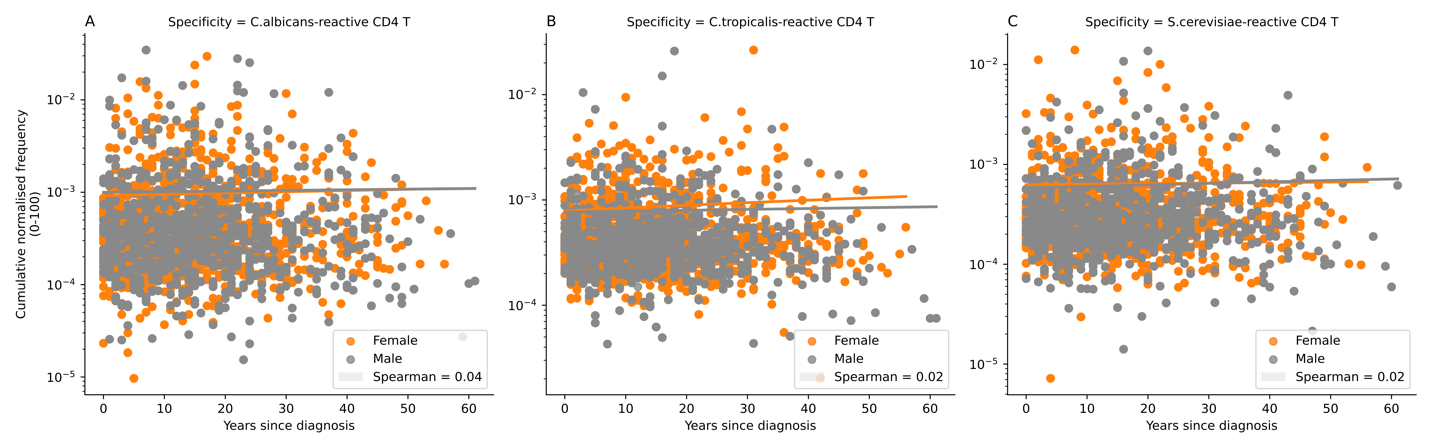

**Figure S3**: The relationship between the expansion of fungal-specific clonotypes and years since diagnosis in individuals from the SPARC IBD cohort.

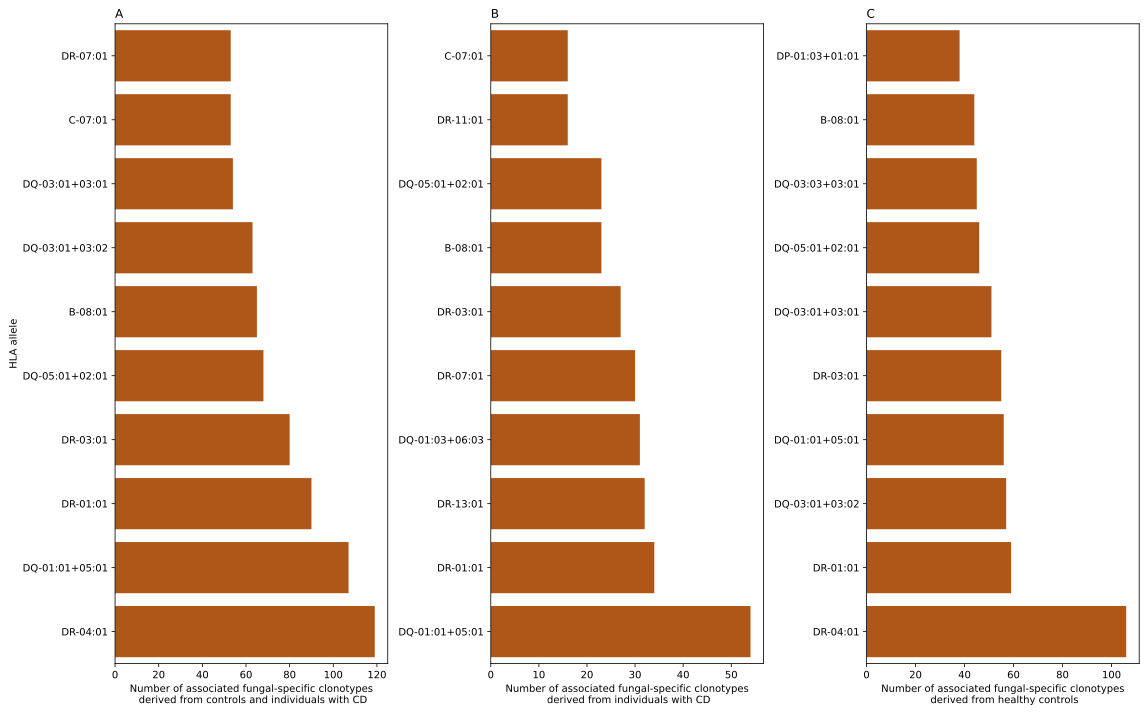

**Figure 4**: the statistically resolved cognate HLA proteins for fungal-specific clonotypes. (**A**) the top-10 alleles that were predicted to interact with fungal specific clonotypes derived from healthy controls and individuals with CD, while (**B**) depicts the HLA restriction map for clonotypes derived from individuals with CD and (**C**) from healthy controls only.

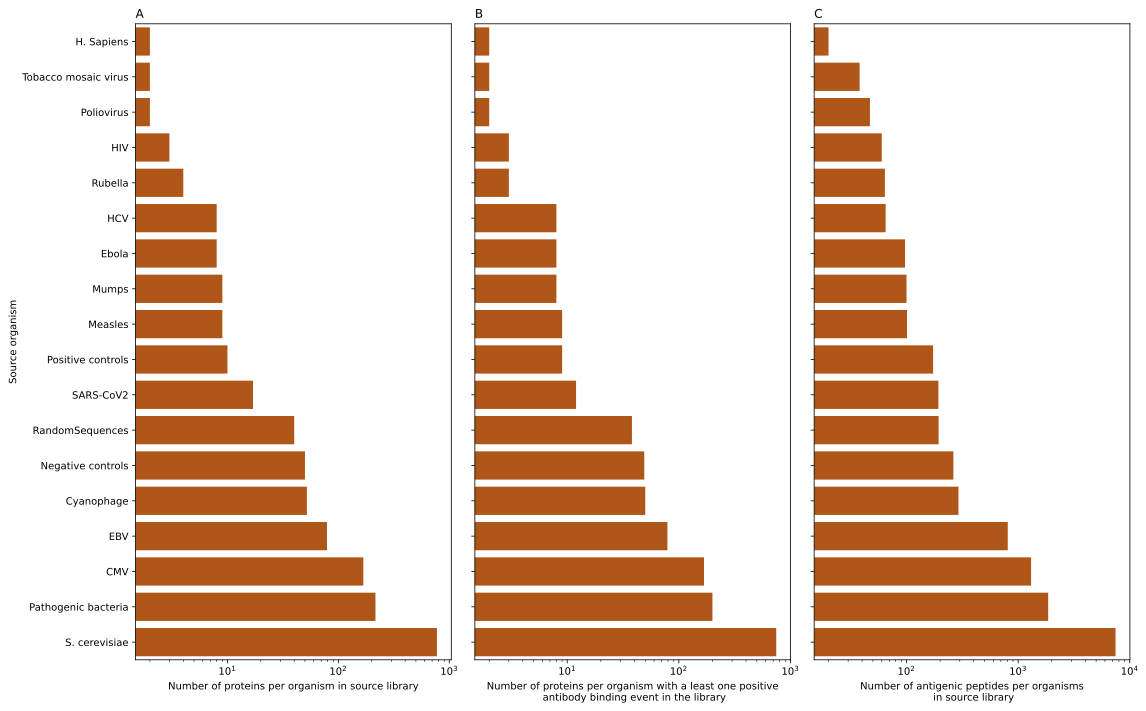

**Figure S5**: the composition of the novel PhIP-Seq library designed to investigate the immunogenicity of S. cerevisiae. (**A**) shows the overall number of proteins included in the final library design from each organism, while (**B**) depicts the overall number of proteins observed after cloning and conducting PhIP-Seq across the 160 samples included in the current study. Lastly, (**C**) depicts the number of antigenic peptides generating a response detected in at least one individual, per organism.

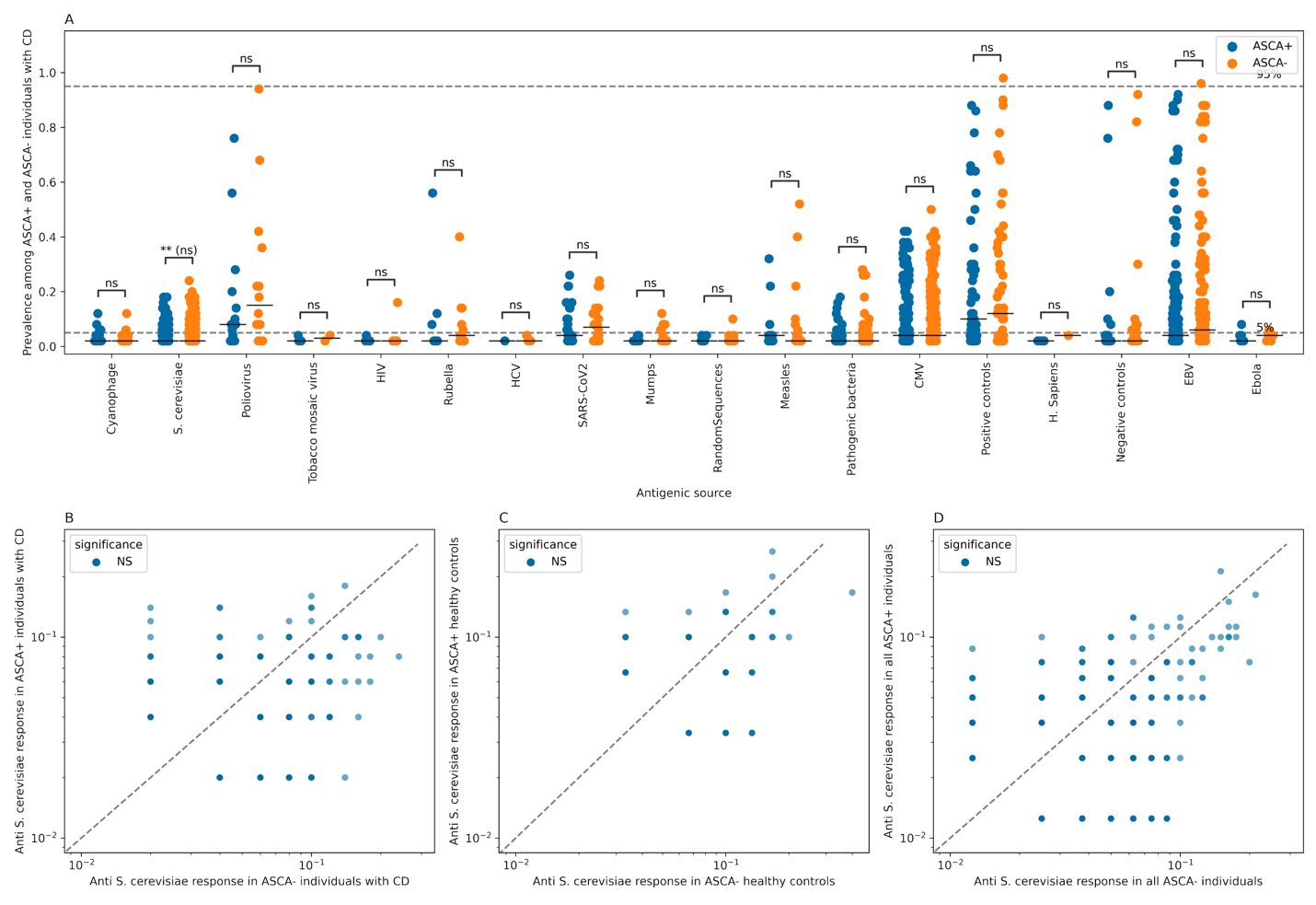

**Figure S6**: ASCA^+^ and ASCA^-^ individuals show a similar response toward S. cerevisiae derived peptide antigen. (**A**) shows a comparison between the prevalence of antibody responses against different antigenic sources in ASCA^+^ and ASCA^-^ individuals with CD. (**B**) compare the prevalence of anti- S. cerevisiae peptides between ASCA^+^ and ASCA^-^ individuals with CD. Lastly, (**C**) shows the difference in response to S. cerevisiae peptides in all ASCA^+^ and ASCA^-^ individuals regardless of their disease status. In **A**, a two-sided Mann-Whitney U test was used to compare the prevalence of immune responses between ASCA^+^ and ASCA^-^ individuals using the Benjamini-Hochberg method for multiple-test correction. In (**B**-**D**), we selected peptides from S. cerevisiae that were detected in both ASCA^+^ and ASCA^-^ individuals and had a difference in prevalence of more than 1%. After that the Fisher’s exact test was used to compare their prevalence in cases and controls, lastly, we used the Benjamini-Hochberg approach to correct for multiple testing.
